# Mutual dependence between membrane phase separation and bacterial division protein dynamics in synthetic cell models

**DOI:** 10.1101/2024.09.17.613417

**Authors:** Nishu Kanwa, Shunshi Kohyama, Leonard Fröhlich, Amogh Desai, Petra Schwille

## Abstract

Cell membranes in bacteria are laterally polarized to produce specific environments for membrane proteins, e.g., proteins involved in cell division which accumulate at mid-cell or the cell poles. An interesting result of such membrane-lipid interplay is the reorganization of lipid domains together with membrane-bound proteins at the onset of cell division, suggesting a functional significance of membrane compartments in the cell cycle. Here, by adopting the key bacterial division proteins MinCDE and FtsZ as an archetypal spatial patterning system, we present a simple vesicle-based *in vitro* model to explore the mutual dependence of protein pattern formation and membrane heterogeneity. Like many other peripheral membrane proteins, MinDE exhibit preferential binding and macro-scale pattern formation at L_d_ domains, which leads to altered oscillation mode selection in phase-separated membrane compartments (GUVs). Moreover, incorporating bacterial division proteins within phase-separated GUVs leads to blebbing-like membrane deformations followed by the reorganization of L_o_ domains aligning at the neck region of the bleb, which agrees well with the domain rearrangement in bacterial membranes immediately preceding the radial constriction process. Overall, the presented *in vitro* model system showcases a basic framework to better comprehend the cellular division mechanism in consideration of complex cellular lipid environments.

## Introduction

With the advanced imaging capabilities of the last decades, it became obvious that like in eukaryotes, bacterial cell membranes demonstrate remarkable characteristics in their spatiotemporal organization. Particularly notable is both spatially and temporally controlled localization of certain lipid species, which generally constitutes a fundamental feature of biological membranes [1]. In the context of bacterial division processes, some topological regions, such as the mid-cell and the cell poles, hold particular significance as they serve as specialized sites for lipid and protein regulation and organization. These regions concentrate essential components that assist in accurately placing and facilitating the function of the divisome – the protein complex responsible for coordinating cell division processes [2].

In fact, the lateral organization of lipids in bacterial cell membranes appears to be characterized by microdomains enriched in specific lipid components, forming sub-micron-sized patches based on their intrinsic chemical properties. These domains act as scaffolds for numerous proteins and are crucial for their recruitment to the membrane surface [3]. As a consequence, protein-enriched lipid domains form within membranes, and such lipid domains are further stabilized and sustained by membrane proteins [4]. The process of phase separation is correlated with the reduction of membrane fluidity *in vivo*. Lipid bilayers primarily exist in the liquid-disordered phase (L_d_) and may separate into nanoscale, hopanoid-containing liquid-ordered phases (L_o_) [3a].

Phase-separated membrane domains are supposed to play a unique role in division processes through regulating functional division machinery. For instance, cardiolipin (CL), a key component of membrane-protein regulation, localizes at the polar and septal regions of *Escherichia coli* [5], maintaining membrane curvature and non-bilayer structures during the different steps of division. At the same time, two other major components of the bacterial membrane - PE and PG - sequester into domains between polar and septal regions [6]. This spatial heterogeneity results in the enrichment of CL in *E. coli* minicells that are derived from the cell poles [5, 7], suggesting that specific lipids might recruit proteins to the poles and the division site in living cells. Indeed, peripheral lipid domains and peripheral proteins appear to co-localize *in vivo*, which underlines the importance of the co-regulation and interplay of both components in living cells. In order to maintain this regulation, lipid domains must continuously reorganize within the cell.

In order to comprehend such intricate protein-membrane dynamics mediated through lipid domains, one of the fundamental challenges has been to reconstitute functional proteins *in vitro*, while taking into account the complexity of the bacterial cell membranes. By adopting reconstituted systems, the spatiotemporal organization of both bacterial proteins and lipids have been individually studied [8], however, their mutual dependence with regard to division process has not been addressed so far. Moreover, although the development of *in vitro* models of eukaryotic cell membranes is relatively well understood [9], the study of membrane compartments in synthetic prokaryotic counterparts has been lacking. While the production of phase-separated GUVs by using cDICE and emulsion transfer methods has been previously reported [10], their one-pot production with biological systems, such as cell-free protein expression systems, has only been demonstrated in more complicated microfluidic setups [11], which calls for an easier methodology to reconstitute proteins within phase-separated GUVs.

Thus, we here aim to study the interplay between membrane domains and protein dynamics by creating a synthetic model system that mimics the key features of bacterial membranes. We generate membrane compartments that exhibit domains resembling those in bacterial cells, by developing an easy-to-use method to reconstitute protein systems in GUVs featuring membrane phase separation. In this model system, we employ bacterial cell division proteins MinCDE, FtsA, and FtsZ, which play a crucial role in targeting the divisome through spatiotemporal membrane binding dynamics. Briefly, MinCDE proteins generate a concentration gradient on the inner membrane that repeatedly oscillates between cell poles via energy-dissipative self-organization dynamics of MinD and MinE. FtsA and FtsZ form ring-like filament structures that lead to early divisome assembly at the mid-cell region as a spatial counterpart of Min protein oscillations, through the negative regulator of FtsZ assembly, MinC. This precise but robust spatial regulation of bacterial cell division has been extensively studied in model *in vitro* systems, but so far not been addressed in combination with membrane phase separation.

Here, we find that in the presence of membrane domains, both Fts filaments and Min proteins show selective binding to the L_d_ domains, consequently skipping the L_o_ domains of the membrane. Moreover, since Min waves are found to be temperature dependent with respect to wave patterns and oscillation period, Min protein dynamics can be regulated via membrane phase separation, which is strongly dependent on temperature. Even more remarkable is the effect of collective divisome protein dynamics on the domains themselves. Co-reconstitution of Min and Fts proteins successfully demonstrates the assembly of Z-ring structures and membrane deformation driven by their constriction forces within phase-separated GUVs. Interestingly, with such higher-order protein structures, lipid domain reorganization is observed, where the Lo domains spontaneously localize at the neck region of the deformed GUVs. Notably, such domain reorganization dynamics are commonly observed *in vivo* during the division process [7b], confirming the physiological significance of the reconstituted system. Thus, the experimental insights reported here demonstrate not only the temperature-dependent activity of Min dynamics, but also the protein-aided reorganization of domains in the deformed vesicles. Overall, this study showcases a complex yet controllable synthetic cell model that mimics bacterial systems with respect to cellular membrane properties. This model serves as a powerful tool for the biophysical characterization of fundamental biological processes that give rise to spatiotemporal protein organization within phase-separated membrane domains. Moreover, our model would pave the way for the design and engineering of synthetic cells with increased complexity, capable of emulating and manipulating these cellular processes *in vitro* environments.

## Results and discussion

### Preferential binding of proteins with phase-separated membranes

We first explore the membrane binding and pattern formation of MinDE proteins on phase-separated membranes. The bacterial Min system forms spatiotemporal patterns on lipid membranes upon sequential cycles of membrane binding and unbinding of MinDE proteins: i) MinD cooperatively binds to the membrane via ATP-dependent dimerization, ii) MinE dimers are recruited by membrane-bound MinD and form a MinDE complex, iii) the MinDE complex induces ATP hydrolysis of MinD, resulting in dissociation of MinDE proteins from the membrane [12]. Importantly, both MinDE proteins bind the membrane via MTS region in their N-or C-terminus, where their affinity is increased by the negative charge content of the lipid surface. Thus, we employed a lipid composition for phase-separated membranes (mixture of DOPC, DOPG, DPPC, DPPG, and cholesterol) in both SLBs and GUVs, resulting in coexisting L_d_ and L_o_ domains with 30% negative charge content to facilitate MinDE binding (Fig. 1a). Moreover, unlike standard electroformation methods for phase-separated GUVs production, we customized a double-emulsion transfer method, allowing efficient encapsulation of proteins inside GUVs while retaining the temperature-dependent phase separation behavior of lipid domains (see methods).

**Figure 1.**
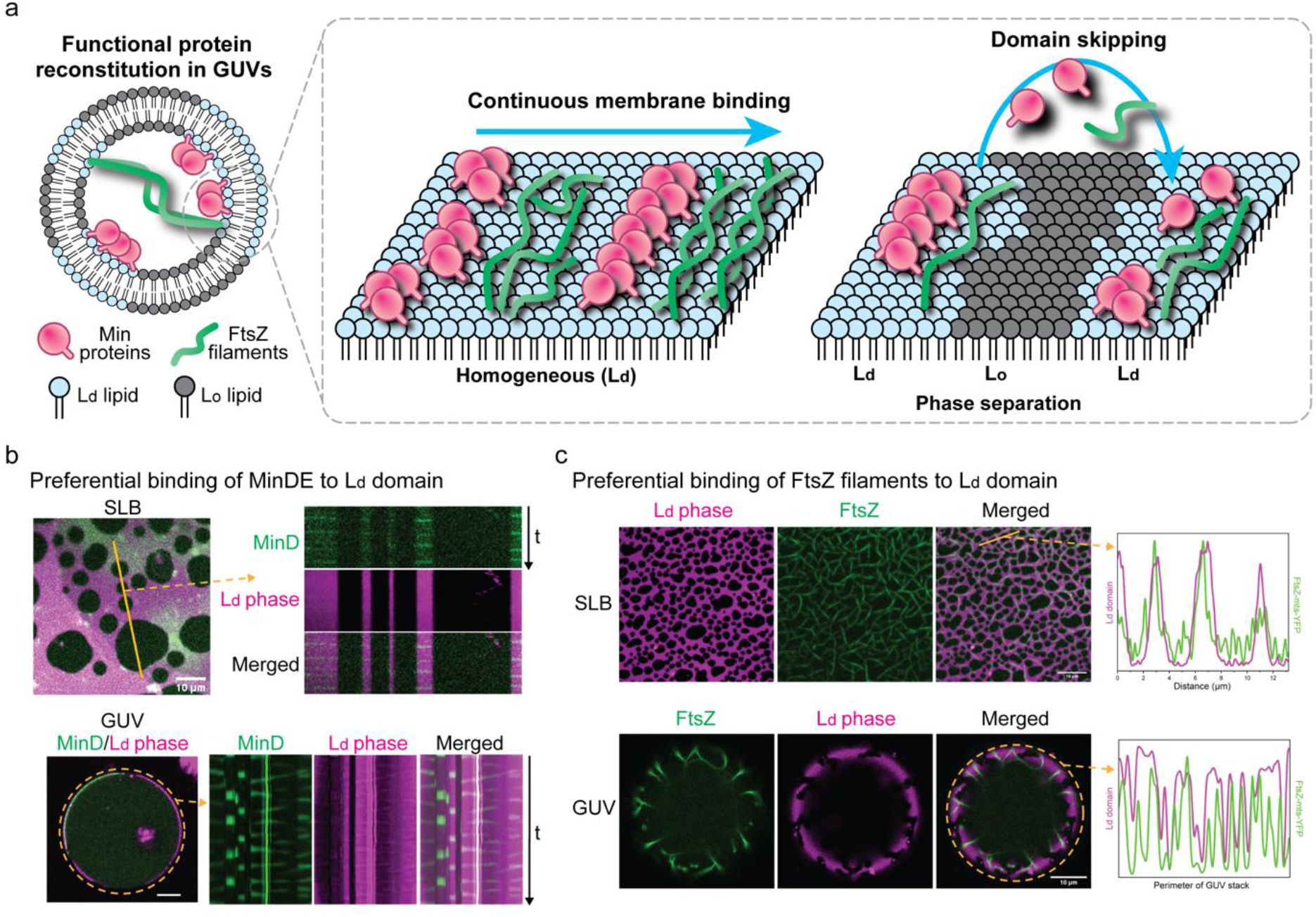
Preferential phase selection dynamics of bacterial division proteins on phase-separated membranes (a) Schematic representation of functional reconstitution of bacterial division protein systems within phase-separated GUVs as well as SLBs consisting of L_d_/L_o_ domains. MinDE and/or FtsZ proteins are encapsulated within GUVs via the transfer-emulsion method. Both proteins preferentially bind to the L_d_ domains, resulting in L_o_ domain skipping on phase-separated membranes, as opposed to continuous binding on homogeneous membranes. Preferential membrane binding of MinDE (b) and FtsZ-mts (c) to L_d_ domain (magenta) of the membrane.

By reconstituting the MinDE protein system on these membranes at room temperature where phase-separated domains emerge, we observed a strong preferential binding of Min proteins for L_d_ domains, resulting in an apparent wave propagation on the L_d_ domain by skipping the L_o_ domains (Fig. 1a-b). Furthermore, since the negative charge of the L_d_ domain serves to be an essential factor in promoting the localization of Min proteins, it must be noted that we did not observe any binding of Min proteins with the membrane in the absence of any negatively charged lipids in the mixture (Fig. S1). However, the negatively charged lipids in L_o_ domains do not facilitate membrane binding of Min proteins suggesting that the binding efficacy is promoted by higher fluidity of the membrane. In order to rule out the possibility of non-incorporation of negatively charged ordered lipid (DPPG) in the L_o_ domains in phase-separated membranes, we observed the interaction of proteins with SLBs composed solely of ordered lipid with the standard 30 % negative charge (DPPC: DPPG, 70:30) and found that the proteins did not show any interaction with the membrane of this composition (Fig. S2), which further indicates that such binding is driven by the high fluidity of the membrane.

Although such selectivity is not surprising, since membrane proteins are commonly observed to preferentially bind to L_d_ domains *in vitro*, our results suggest that the membrane charge efficacy in protein-membrane interactions can be excluded in the L_o_ phase. This corroborates with the strong evidence of the spatially distinct localization of highly fluid membrane microdomains *in vivo*, and co-localization of bacterial membrane proteins with fluid lipid domains. This *in vitro* lipid phase-driven protein organization may account for the recruitment of septal and polar localization of division-related proteins *in vivo*.

Next, we observed the membrane binding preference of FtsZ filaments and found a similar trend, in that FtsZ filaments localize preferentially on the L_d_ domains (Fig 1a, c). Analysis of intensity spectra clearly reveals that the FtsZ filaments avoid the L_o_ domains in both SLBs and GUVs (Fig 1c), demonstrating the significance of membrane domains in regulating the specificity of FtsZ membrane binding as well as MinDE. Following the above observations, we conclude that the membrane can be experimentally compartmentalized into phase-separated domains, where the L_d_ domain is functional for protein localization, but the L_o_ domain is not. Even though the lipid composition in our system is different from that found in bacteria, we observe a similar trend for membrane-protein binding, where the division proteins segregate with lipid domains based on their high fluidity [13].

### Oscillation mode selection of Min proteins in phase-separated GUVs

Intracellular patterns of Min waves, in particular pole-to-pole oscillations, are known to play an important role in the positioning of cell division protein assembly. However, in the absence of external cues such as cell shape, co-supplemented molecular objects on membranes, or lipid heterogeneity in the membrane, Min patterns are mostly random and there is no predominant wave pattern within GUV-based reconstituted systems [8b]. Since lipid heterogeneity shows preferential binding of Min proteins, we explored the possibility of breaking this non-selectivity, resulting in preferential patterns as well. We examine this feature as a function of temperature by utilizing the Peltier device-based precise temperature control in phase-separated GUVs as a cell-mimetic model.

Generally, three major dynamic Min oscillation modes inside micron-sized compartments are found (Fig. 2a), i.e., pulsing (homogeneous cycling between bulk and membrane of the GUV), pole-to-pole oscillations (spatially bi-stable oscillation between two poles of the GUV membrane), and traveling waves (directional propagation of protein waves along membranes). Inside the homogeneous membrane system, the majority (72%) of Min wave patterns were originally found to be pulsing at room temperature, similar to what we previously quantitated [8b], and the rest of the pattern indicates traveling waves. However, strikingly, the wave patterns were completely dominated by pulsing once the temperature was increased to 37 °C, and the pulsing pattern was maintained afterward regardless of the temperature until 22 °C at least (Fig. 2b). Importantly, this is a clear contradiction to previous reports, where reconstituted Min waves showed strong hysteresis characteristics both on SLBs and in microdroplets, after a change of experimental parameters, such as MinDE ratio and temperature [14].

**Figure 2.**
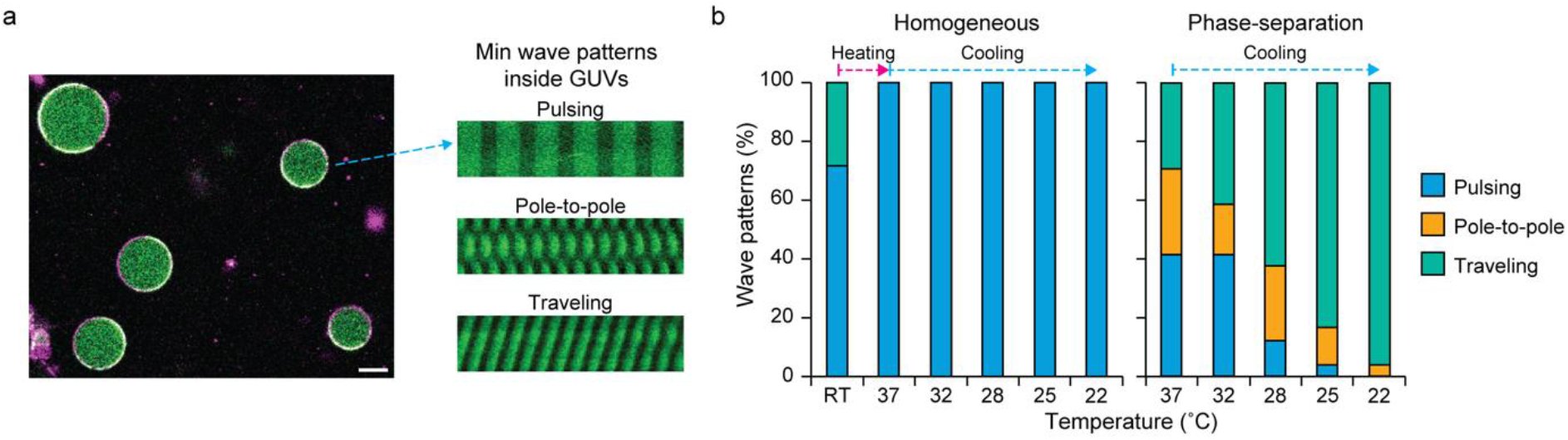
Temperature dependency of Min wave patterns in phase-separated GUVs. (a) A representative snapshot of phase-separated GUVs (L_d_ phases are indicated in magenta) containing mGL-MinD (indicated in green) and MinE proteins and kymographs of three major Min oscillation modes within GUVs. Scale bar: 20 μm. (b) Mode distribution within phase-separated or homogeneous membranes shows a strong temperature dependence of wave patterns within phase-separated membranes (right), whereas only the pulsing mode was found within homogeneous membranes regardless the temperature after once it reaches 37 °C. suggesting that membrane heterogeneity affects the preferential membrane binding of Min proteins.

This discrepancy called for further quantification of the system, where we found a strong temperature dependence of Min oscillation modes within phase-separated GUVs, in stark contrast to the homogeneous membrane (Fig. 2b). At higher temperatures (37°C), Min dynamics show all three oscillation modes at relatively equal probability (pulsing: 42%, pole-to-pole: 29%, and traveling: 29%). However, with decreasing temperatures, traveling waves gradually dominate the pattern distribution, resulting in a reduction of pulsing and pole-to-pole modes among the population. Eventually, the pulsing patterns, which are exclusive in homogeneous membranes, completely vanish at 22°C, and the traveling waves become predominant (96%). This indicates that local heterogeneity of the membrane composition, which strongly affects the preferential membrane binding of Min proteins, also results in a strong mode selection of Min wave. Also, it is worth noticing that we did not observe any pole-to-pole oscillation modes with homogeneous membranes (as in good agreement with that only 1.8% population of this mode was observed in a similar experimental setup with homogeneous membranes in our previous study [8b]), whereas the highest share (29%) of this mode was found with phase-separated membrane at physiological temperature (37 °C). This adds further evidence that bacterial cells utilize membrane heterogeneity to maintain pole-to-pole oscillations by accumulating specific types of lipids, such as CL, at the cell poles.

### Temperature dependency of MinDE oscillation period and reaction kinetics

One of the most crucial parameters for membrane phase separation is temperature. Although most *in vivo* environments exhibit highly mixed lipid composition, which makes it difficult to comprehend the phase-separating behaviors, clear temperature dependency of phase separation is often easily observed *in vitro* environments. Importantly, MinDE oscillation in *E. coli* cells is also known to be strongly temperature-dependent; in particular, it can be described as Arrhenius temperature dependence, in which the enzymatic turnover by MinD (as an overall determinant of oscillation period) increases with temperature according to the Arrhenius law (k = Ae^−Ea/RT^). Therefore, it is crucial to study how such temperature-dependent phase behavior affects MinDE oscillations in controlled environments. Thus, we analyzed the temperature dependence of MinDE oscillations based on two essential parameters as known from *in vivo* Min dynamics: (i) oscillation periods and (ii) reaction kinetics.

We quantified the Min oscillation periods, first with homogeneous lipid composition as a control. We found that MinDE oscillations gradually become slower, with longer molecular residence times on the membrane and higher amplitudes at lower temperatures (oscillation period of 120.1 sec at 22 °C) as compared to physiological temperatures (38.3 sec at 37 °C), showing roughly 3 times slower oscillations over 15°C change in temperature (Fig. 3ab, movie S1). A similar trend can be found for phase-separated vesicles. However, with decreasing temperature, significant differences (p ≤ 0.0001) are observed in the oscillation period at temperatures lower than 28 °C, where L_o_ domains start originating (Fig. 3a, indicated by yellow triangles), resulting in phase-separation in the membrane. Eventually, at 22 °C, the average oscillation period further slows down to 177.0 sec, roughly 4.5 times slower than 37 °C (Fig. 3c). As we found no significant difference (p = 0.62) in the oscillation period at physiological temperature (39.1 sec at 37 °C in phase-separated GUVs) compared to the homogeneous membrane (38.3 sec) (Fig. 3c), we concluded that phase-separated domains also change the reaction kinetics. To support this, among phase-separated GUVs, there is a weak dependence (r = 0.28, Fig. 3d) between the occupancy of L_d_ phases in the membrane and Min oscillation period, suggesting that L_d_/L_o_ appearance is a contributing factor to the wave dynamics, where oscillation periods become slower with an increase in L_d_ area coverage upon emergence of phase separation.

**Figure 3.**
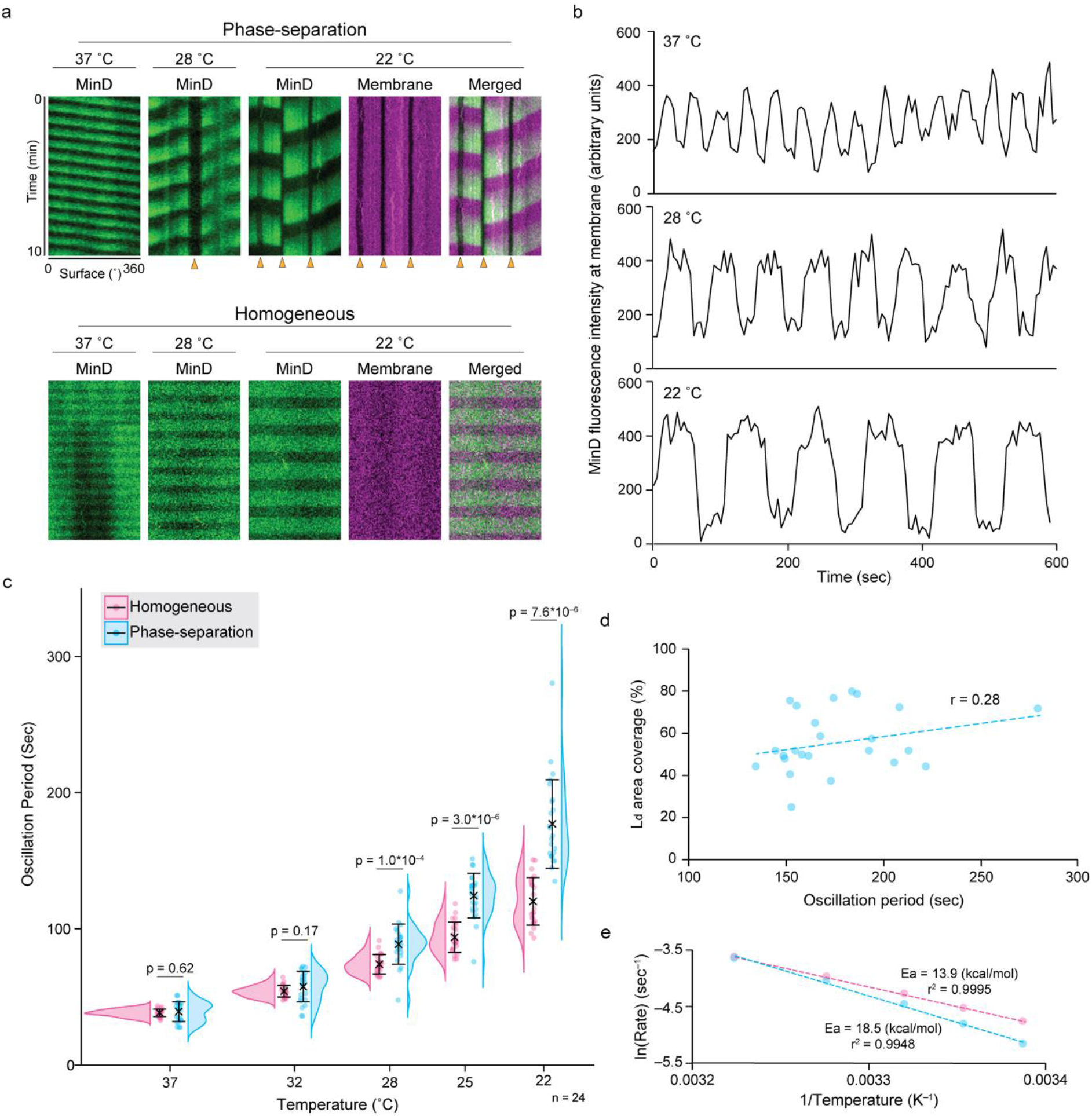
Effect of temperature on MinDE dynamics. (a) Kymographs of MinD oscillations within GUVs composed of phase-separation or homogeneous membrane. In both phase-separated and homogeneous membrane, Min oscillations slow down with decreasing temperature. L_o_ domains (indicated by yellow triangles) appear in phase-separating membrane below 28 °C, inducing preferential MinD binding on L_d_ phases, while homogeneous membrane shows no phase separation even at 22 °C. (b) MinDE oscillation dynamics (mGL-MinD fluorescence intensity on the membrane) at different temperatures. Slower oscillation periods, longer residence times on the membrane, and higher amplitudes can be found at lower temperatures. (c) Violin plots of the MinDE oscillation periods within GUVs. Periods are significantly slowed down below 28 °C. Plots indicate all the data points for analysis. Black lines and cross symbols indicate average and standard deviations. (d) Scatter plot of L_d_ area coverage vs. Min oscillation period within phase-separated GUVs shows a weak dependence (r = 0.28), suggesting the reactivity of MinDE on the membrane may slow down with the higher L_d_ coverage. (e) Arrhenius plot of the temperature-dependent reaction rate of Min proteins within GUVs. Phase-separation membrane shows higher temperature dependence with activation energy (E_a_) of 18.5 kcal/mol, while homogeneous membranes show lower E_a_ of 13.9 kcal/mol.

Furthermore, the Arrhenius plot (Fig. 3e) confirms the apparent difference in the activation energy (E_a_) for Min oscillations, indicating higher E_a_ on the phase-separated membrane (18.5 kcal/mol) compared to the homogeneous membrane (13.9 kcal/mol). This indicates that the reaction rate of Min proteins is much higher within the homogeneous membrane. However, strikingly, we found the higher E_a_ with the phase-separated membrane is in good agreement with the previously measured activation energy (20 kcal/mol) in *E. coli* cells. Together with the similar magnitude of change in the oscillation period (roughly 4.5 times in this study compared to 4 times in *E. coli* cells (between 20 °C and 40 °C in [15])) as well as the temperature-dependent growth rate of *E. coli* cells (4 times difference between 21 °C and 37 °C), which is also known to follow the Arrhenius dependence, our results correspond well with observations *in vivo*, suggesting the cellular membrane environment to be rather heterogeneous. Thus, our results indicate that membrane phase separation plays a key role in controlling the cellular Min oscillation period that eventually contributes to the cell growth rate in different temperature environments, highlighting the relevance of membrane heterogeneity with regard to protein dynamics in bacterial cells.

### Alignment of FtsZ filaments at mid-cell triggered by temperature

Another hallmark of the *in vivo* Min system is its positioning of the division ring based on FtsZ protein, the so-called Z-ring, to mid-cell. As demonstrated previously, both Min and FtsZ dynamics are sensitive to membrane phases and exhibit preferential binding to L_d_ domains. Thus, here we co-reconstitute both proteins and explore the assembly and positioning of Z-ring structures inside phase-separated GUVs, investigating how membrane phase separation behavior influences the emergence of such a higher-order protein structure. We have previously demonstrated the co-reconstitution of MinCDE and FtsZ inside homogeneous GUVs [8b], allowing to study Z ring assembly systems *in vitro* environments. Here we adopted the co-reconstitution assay for MinCDE/FtsZ protein system inside phase-separated GUVs, where the membrane domains could be mixed and de-mixed by temperature cycling (Fig 4a).

**Figure 4.**
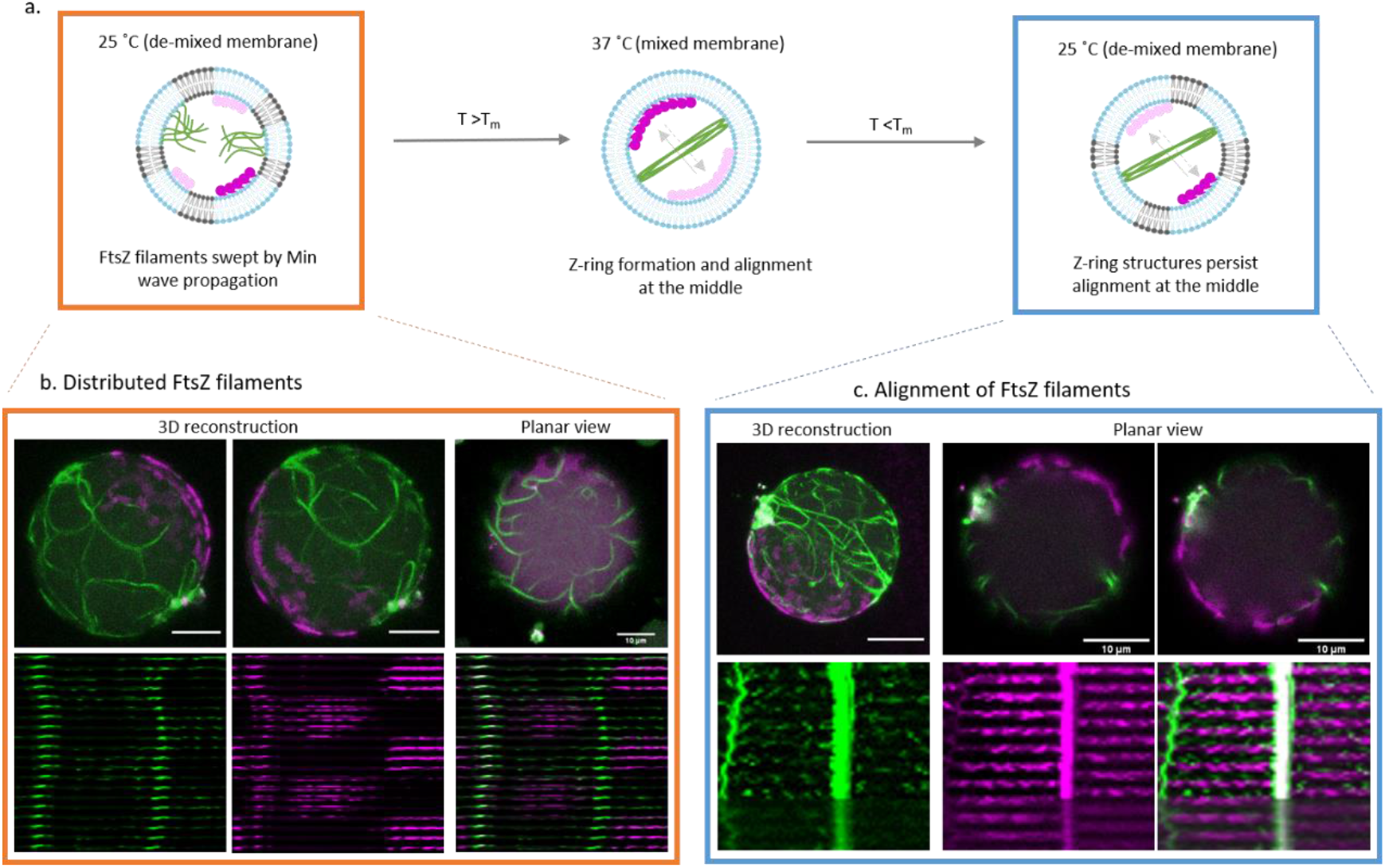
Min/FtsZ co-reconstitution confirmed FtsZ-ring formation within phase-separated GUVs. Z-ring formation is robust over the mixing/demixing processes and becomes more prominent by temperature cycling (a), i.e. temperature ramps ranging from 25°C to 37°C. Both conditions depicted here represent protein dynamics in the presence of domains at room temperature (25°C) before (b) and after (c) the domain mixing cycle. Scale bar: 10 μm.

As demonstrated, both FtsZ filaments and MinCDE waves appear at L_d_ domains, evidently skipping micron-sized L_o_ domains in the GUVs (movie S2). However, unlike in homogeneous membranes [8b], the FtsZ-ring structure does initially not form in phase-separated GUVs at room temperature (RT, below T_m_) (Fig. 4b). This could be attributed to the prevalently observed traveling Min patterns at this temperature (Fig. 2a-b as discussed above), where as a consequence, the FtsZ filaments spread out over the entire L_d_ domain, and are continuously swept away by the traveling Min waves, rather than condensed into a ring structure in the MinDE oscillation minima (movie S2).

Strikingly, when we increased the temperature above T_m_ (until 37 °C) to induce the mixing of domains, MinCDE patterns developed over the entire fluid membrane surface, resulting in an increased probability of pole-to-pole oscillation mode (Fig. 2b), which allowed Min proteins to position FtsZ filaments into ring-like structures (Fig 4c). Remarkably, once the Z-ring structures are positioned in the middle of GUVs, the ring structures remain stable during the subsequent cooling from 37°C to below T_m_ (observation at 25°C) and even after the re-emergence of micron-sized L_o_ domains. Moreover, during the temperature-induced de-mixing in the presence of Z-ring structures, micron-sized L_o_ domains start appearing in the protein-free membrane regions, while FtsZ filaments stabilize L_d_ domains underneath, due to their higher L_d_ affinity. Similar reorganization of lipid domains controlled by the pre-existing protein filament networks has already been reported for the actin networks on SLBs [16], indicating that protein assemblies can spatiotemporally control the local lipid phase during phase transition, e.g. due to temperature change (movie S3). Taken together, we not only found the critical conditions to assemble Z-ring structures in phase-separated GUVs guided by Min waves, but we could implement additional control of protein filament assembly in our synthetic cell models by employing temperature as a trigger, suggesting a novel strategy for desired synthetic cell deformation.

### Activation of Min/FtsZ dynamics in phase-separated GUVs via a cell-free expression system

To further examine the functionality of reconstituted biological systems inside phase-separated GUVs, we employed a cell-free protein expression system, which is not regularly used in such complex membrane systems. The only example so far has been the expression of GFP inside microfluidically generated phase-separated GUVs [11]. However, the full potential of cell-free expression in cell models with more complex membranes, including the *situ* synthesis of functional membrane-interacting proteins, has not been demonstrated yet. As we previously have shown that MinCDE, FtsZ, and FtsA proteins can be produced via the cell-free expression system PURE [17] inside homogeneous GUVs [8b], we conducted protein production by PURE in phase-separated GUVs generated by the adapted double-emulsion transfer method.

We first performed the PURE cell-free expression of either MinCDE or FtsAZ proteins as sub-systems of bacterial early divisome assembly. We confirmed that both systems were successfully reconstituted within phase-separated GUVs by expressing those proteins at 37 °C (above T_m_: note that at this temperature, membrane domains are mixed) as the working temperature of PURE system (Fig. 5a, S3a, movie S4, S5). Moreover, by lowering the temperature below the T_m_ after the successful expression of these proteins, phase separation was initiated, resulting in the partitioning of MinCDE to the emerging L_d_ domains. We also confirmed that the oscillation period slows down with a decrease in temperature (Fig. 5a, movie S4), confirming the temperature and phase separation-dependent oscillation control shown in Fig. 3. In the case of FtsZ and FtsA expression, we found the same spatial appearance of L_o_ domains as reported above, i.e., micron-sized L_o_ domains avoid FtsZ filaments and appear only at the protein-free regions during the de-mixing. Thus, based on these observations, we concluded that we could successfully functionalize PURE cell-free expression inside phase-separated GUVs, providing an easy-to-use methodology for bottom-up construction of more complex cell models *in vitro*.

**Figure 5.**
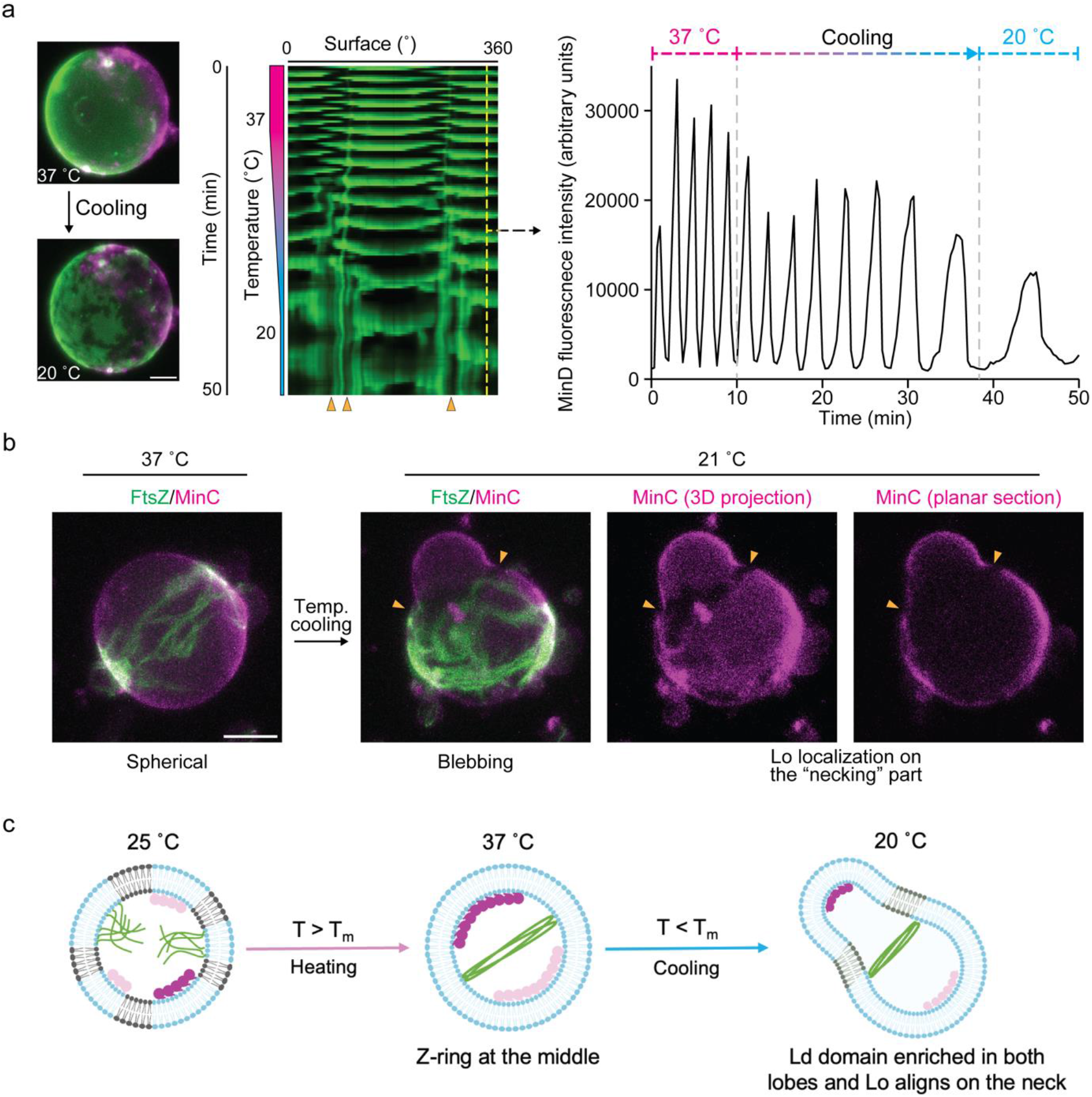
Reconstituted Min oscillation and FtsZ-ring formation in phase-separated GUVs via PURE cell-free system. (a) Representative images and kymograph of oscillation dynamics of cell-free synthesized MinDE proteins within phase-separated GUVs (msfGFP-MinC is indicated in green). Min dynamics with an intensity profile shows that Min oscillations slow down with decreasing temperature. Also, L_o_ regions (indicated by yellow triangles) emerge at lower temperatures. Scale bar: 10 μm. (b) Membrane blebbing of a phase-separated GUV containing FtsZ-ring structures. The vesicle first remains at a spherical shape at 37 °C after the cell-free expression of MinCDE (mCherry2-MinC is indicated in magenta) and FtsAZ (FtsZ-G55-Venus-Q56 is indicated in green) proteins. After cooling, a part of the membrane starts blebbing, with reorganization of the L_o_ phase at the low-curvature neck of the bleb (indicated by yellow triangles). Scale bar: 10 μm. (c) Schematic representation of MinCDE and FtsZ filaments’ antagonistic placement, alignment of FtsZ at mid-cell triggered by temperature and pole-to-pole patterns of Min proteins, followed by vesicle deformation and re-alignment of domains.

### Blebbing and rearrangement of membrane domains in phase-separated GUVs constricted by a minimal Z-ring

Finally, we studied how the membrane phase separation behavior affects the Z-ring constriction of GUVs, aiming to fundamentally understand the interplay between the bacterial division machinery and cellular membranes retaining phase-separated domains. Following the reconstitution of sub-systems, we performed co-expression of MinCDE, FtsZ, and FtsA within phase-separated GUVs, allowing the emergence of FtsZ-ring formation at 37°C, followed by segregation of membrane into phase-separated domains arising with the decrease in temperature from ∼28°C (Fig. S3b, movie S6). The following observations were made: (i) randomly distributed FtsZ filaments develop into a meshwork on the membrane, (ii) FtsZ meshes reorganize into highly condensed, ring-like structures guided by Min oscillations, (iii) the phase-separated membrane domains emerge via temperature cooling while keeping FtsZ-ring structures in position, as already observed in both purified protein or cell-free expression coupled systems.

Strikingly, we confirmed the large-scale deformation of the GUVs into membrane blebs by following the Z-ring driven membrane constriction (Fig. 5b, movie S7). Initially, the GUVs maintain a rather spherical shape at 37 °C with the Z-ring positioned in the middle, however, after the cooling process, a part of the GUV membrane starts blebbing, with a stable localization of L_o_ phase at the neck of the bleb, as depicted in Fig 5b-c. This is accompanied by L_d_ domains aligning with the two lobes of the deformed GUV, while L_o_ remains at the neck of the deformed GUV, thereby remaining in a low curvature region. Such membrane curvature-driven lipid sorting has previously been observed where L_o_ domains in phase-separated membranes align at regions of low curvature [18]. Moreover, lipid sorting in CL containing GUVs has been demonstrated by following membrane tube pulling assays [19].

Importantly, the Z-ring structure remains stabilized in the middle of the GUV and is not shifted to the neck region, suggesting that phase separation plays a critical role in blebbing in this scenario. While the placement of the L_o_ domain on less-curved regions could be explained by a decrease in line tension [20], such a large-scale shape deformation under an iso-osmotic condition likely results from the force imparted by Min dynamics in combination with the Z-ring. In particular, pole-to-pole Min oscillations may induce deformation in the GUV to produce the bleb in the GUVs, while Z-ring helps with the stabilization of this deformed shape, subsequently allowing the reorganization of lipid domains in order to localize L_o_ domains at the neck. Since such reorganization of lipid domains is commonly observed *in vivo* environments during cell division in the *E. coli* membranes [7b], we here provide an important case study in simplified *in vitro* setups of how complex spatiotemporal protein dynamics emerge together with phase-separated membrane dynamics. Our results not only demonstrate the successful reconstitution of cell-free expression-assisted assembly of division machinery, but also point to a potential role of lipid phase separation in bacterial cell division.

## Conclusion

In the present study, we successfully developed a one-pot GUV-based synthetic cell system produced by an easy-to-use double-emulsion transfer method exhibiting membrane heterogeneity with tunable charge. We examine the intricate interplay of lipid heterogeneity and bacterial division proteins (Min/Fts) dynamics and observe the preferential protein binding to L_d_ domains. The wave dynamics of Min proteins are found to be temperature dependent, as also observed *in vivo*, following the Arrhenius law. Although it is not surprising that Min waves are found to be temperature dependent, the activation energy given by the Arrhenius equation shows a stark difference between homogeneous and phase-separated membranes, in that phase-separated membranes more closely resemble the *in vivo* dynamics of Min proteins. Interestingly, the interplay between MinDE oscillations and Z-ring alignment deforms the GUVs to form membrane blebs, followed by a rearrangement of the membrane domains. This appears to correspond to the domain reorganization found *in vivo* during the division process. Taken together, membrane heterogeneity serves as an important parameter and must be further explored to comprehend cellular processes which require a large-scale interplay between membrane domains and membrane-associated proteins. On the other hand, based on presented technical advancements, membrane heterogeneity is a fascinating feature for developing more sophisticated *in vitro* cell models. In combination with other synthetic biology tools, such as dedicated microfluidic devices or light-controllable azo-lipids, the introduction of more complex membrane composition might offer an additional layer of functionality, e.g., for enhanced membrane shape deformations within GUVs.

## Acknowledgments

The authors would like to thank the MPIB Core Facility, Michaela Schaper, Katharina Nakel, Kerstin Röhrl, Beatrix Scheffer, Sandra Ortmeier and Sigrid Bauer for assistance in protein purification, plasmid construction and lipid preparation. The authors thank Maria Reverte Lopez for useful scientific advice and helpful comments. We also thank Dr. Kareem Al Nahas and Dr. Henri Franquelim for their valuable feedback and discussions. This work is supported by the Deutsche Forschungsgemeinschaft (PS). SK is supported by JSPS Overseas Research Fellowships. The authors would also like to acknowledge the support of the Center for Nanoscience (CeNS), Munich.

