## Supplementary figure 1-3, Methods for "Mutual dependence between membrane phase separation and bacterial division protein dynamics in synthetic cell models"

### Supplementary Information

#### Materials and methods

*Plasmid construction:*

All DNA fragments were amplified by using Phusion High-Fidelity DNA Polymerase (Thermo Fisher Scientific, Waltham, MA, USA) and treated with DpnI (Thermo Fisher Scientific). Then, fragments were assembled using GeneArt Seamless Cloning and Assembly Enzyme Mix (Thermo Fisher Scientific). In the case of deleting gene sequences, blunt-end cloning was used to phosphorylate an amplified DNA fragment by T4 Phosphokinase (Thermo Fisher Scientific) and then ligate with T4 DNA Ligase (Thermo Fisher Scientific).

To construct pET28a-mGreenLantern-MinD, the plasmid backbone together with MinD sequence amplified from pET28a-EGFP-MinD [1] with a primer set (gaattcgcacgcattattgttg and ggatccgcgacccatttg) and mGreenLantern [2] gene amplified from a gBlocks gene fragment (Integrated DNA Technologies) with a primer set (gaattctttgtagagctcatc and tcgcggatccatggttagtaaaggagaagaatta) were assembled using seamless cloning.

pPT1-FtsZopt-G55-Venus-Q56 was obtained by transferring FtsZopt-G55-Venus-Q56 gene amplified from pET11b-FtsZopt-G55-Venus-Q56 [3] with a primer set (gttatgctagttaatcagcttgcttacgcag and tttgtttaactttaagaaggagatataccatg) into pPT1 backbones amplified from pPT1-FtsAopt [3] amplified with primer sets (ctagcataaccccttggg and catggtatatctccttcttaaagttaa) and (tgaaacatggcaaaggtagcgt and gctacctttgccatgtttcagaaa) using seamless cloning.

To obtain pPT1-mCherry2-MinC, pET28a-His-mCherry-MinC [3] was first optimized by removing His-tag using blunt-end cloning with a primer set (catggtatatctccttcttaaagttaaac and gtgagcaagggcgaggag) (pET28a-oHis-mCherry-MinC). Then, mCherry sequence was further optimized for cell-free expression by introducing mutations in N-termiuns sequence using seamless cloning with primer sets (gatataccatggttagtaaaggtgaagaagataatatggccatcatcaag and cttgatgatggccatattatcttcttcacctttactaaccatggtatatc) and (tgaaacatggcaaaggtagcgt and gctacctttgccatgtttcagaaa) (pET28a-oHis-mCherryopt-MinC). mCherryopt-MinC sequence amplified with a primer set (gggttatgctaggatcctcaatttaacggttgaacggtc and tttgtttaactttaagaaggagatataccatg) was then transferred into pPT1 backbones (obtained from pPT1-FtsAopt as described above) by using seamless cloning (pPT1-mCerryopt-MinC). Lastly, pPT1 backbones together with MinC gene amplified with primer sets (accttgaagcgcatgaactc and gctacctttgccatgtttcagaaa) or (gcatggacgagctgtacaag and tgaaacatggcaaaggtagcgt) were combined with mCherry2 gene amplified from gBlock gene fragment (Integrated DNA Technologies) with a primer set (gagttcatgcgcttcaaggt and cttgtacagctcgtccatgc).

To construct pPT1-MinD.MinE.T7term, A DNA fragment was amplified from pPT1-MinD.MinE [3] with a primer set (ctagcataaccccttggg and ttatttcagctcttctgcttc), and then ligated using blunt-end cloning to delete redundant sequences between the stop codon of MinE and T7 terminator sequence. All gene sequences were validated using Sanger Sequencing Service (Microsynth AG, Balgach, Switzerland).

*Protein purification:*

Purification of His-MinD [1], His-EGFP-MinD [1], His-MinE [1], MinE-His [1], His-mScarlet-I-MinC [3], His-msfGFP-MinC [3] FtsZ-Venus-mts [4] were descried previously and His-mGreenLantern-MinD (His-mGL-MinD) was obtained according to the purification of His-mScarlet-I-MinC [3]. Briefly, *E. coli* BL21 (DE3) pLysS cells were transformed with pET28-mGreenLantern-MinD and inoculated in LB medium (with 50 μg/mL Kanamycin). The cell culture was then incubated at 37 °C while shaking at 180 rpm until an optical density at 600 nm reached 0.2. Isopropyl-β-D-thiogalactopyranoside was then added at 1 mM to induce protein overexpression, and the cell culture was further incubated for 5h at 37 °C while shaking at 180 rpm. Cells were then pelleted by centrifugation for 3 min at 8,000 × g at 4 °C and stored at –80 ˚C until further purification. The cell pellet was thawed on ice and then resuspended in 3.5 mL Lysis buffer (50 mM Tris-HCl, pH 7.5, 300 mM NaCl, 10 mM Imidazole, 1 mM ADP) and lysed using a tip sonicator (Branson ultrasonics S-250D, Thermo Fisher Scientific). The crude cell lysate was then centrifuged for 30 min at 20,000 × g at 4 °C, and the supernatant was mixed with 1 mL Ni-NTA agarose (QIAGEN, Hilden, Germany) equilibrated with Lysis buffer, gently shaken for 10 min at 4 °C. The Ni-NTA agarose was then loaded into an empty column, washed with 25 mL Wash buffer (50 mM Tris-HCl, pH 7.5, 300 mM NaCl, 20 mM Imidazole, 10% Glycerol), and His-mGL-MinD was eluted with 2 mL Elution buffer (50 mM Tris-HCl, pH 7.5, 300 mM NaCl, 250 mM Imidazole, 10% Glycerol). The buffer was exchanged with Storage buffer (50 mM Tris-HCl, pH 7.5, 150 mM GluK, 5 mM GluMg, 1 mM ADP, 10% Glycerol) using Amicon Ultra-0.5 centrifugal filter unit 30 kDa (Merck KGaA, Darmstadt, Germany). Protein concentration was quantified by using Bradford assay kit (Bio-Rad, Hercules, CA, USA).

*Self-organization assay on SLBs:*

SLBs were prepared via fusion of Small Unilamellar Vesicles (SUVs) deposited on top of freshly cleaved mica already glued on top of a glass coverslip [5]. The lipid mix used is DOPC: DOPG: DPPC: DPPG: Chol (Avanti Polar Lipids, Alabaster, AL, USA) in equal final ratios for phase-separated SLBs labelled with 0.001 mol % Atto 655-DOPE. The lipid mix (4mg/mL in chloroform) was first dried under $N_2$ gas flow, hydrated with Min buffer containing 25 mM Tris-HCl, pH 7.5, 150 mM KCl, 5 mM $MgCl_2$ to yield a final lipid concentration of 0.5 mg/mL, and then sonicated for about 10 min to form SUVs. After the SUV solution became clear by sonication, SUVs (150 µL solution) were added on top of mica and incubated at 50 ̊C for 5 min to form SLBs, which was then rinsed several times with the same buffer. At the end, 300 µL of the sample volume was kept in the chamber before imaging.

For reconstitution of the MinDE system (Fig. 1), His-MinD (0.5 μM), His-EGFP-MinD (0.5 μM), and His-MinE (4 µM) were mixed with 2.5 mM ATP-Mg in Min buffer. For reconstitution of FtsZ (Fig. 1), FtsZ-Venus-mts (0.6 µM) was added to SLBs together with 1 mM GTP-Mg in Min buffer to trigger its polymerization. Confocal images were collected after ~10 min of incubation at RT.

*GUV preparation:*

GUVs were prepared by the inverted double-emulsion transfer method [3] with some changes as described below. The lipid mix used was DOPC: DOPG: DPPC: DPPG: Chol (17.5: 7.5: 31.5: 13.5: 30 mol ratio) labelled with 0.001 mol% Atto655-DOPE to label the Ld phase. As a control, the mixture of POPC: POPG (70:30 mol ratio) labelled with 0.1 mol% Atto655-DOPE was used. The lipid mix (32 mM, 50 µL) was dissolved in chloroform and dried in a glass vial under $N_2$ gas flow for ~15 minutes. The dried film was then suspended in a mixture of decane (20 µL) and mineral oil (500 µL) to yield the final concentration of 3.2 mM and sonicated at elevated temperatures (~50 ˚C) for ~30 minutes. The resultant lipid-oil mix (200 uL) was placed on top of 500 µL of Reaction buffer (50 mM Tris-HCl, pH 7.5, 150 mM GluK, 5 mM GluMg) in a 1.5 mL tube. Another 250 uL of lipid-oil mix (preincubated at 37 ̊C) was mixed with 5uL of inner solution (see next sections for composition) in another tube and emulsified by tapping. 200 µL of this emulsion was then placed on top of the multilayered Reaction buffer/lipid-oil mix solution. This tube was centrifuged at 6000 rcf for 30 minutes at 37 ̊C. The sample was allowed to cool to room temperature for nearly 30 minutes before imaging. The GUVs were then collected from the Reaction buffer and placed on the imaging chambers.

*Encapsulation of purified proteins (MinCDE and/or FtsZ-mts) inside vesicles:*

All inner solutions for purified proteins were prepared in the Reaction buffer. For reconstitution of the MinDE system (Fig. 1), an inner solution containing His-MinD (1.5 µM), His-mGL-MinD (1.5 µM), and MinE-His (3 µM) was mixed with 30 g/L BSA and 2.5 mM ATP-Mg. For the analysis of temperature dependency of Min wave dynamics (Fig. 2-3), His-mGL-MinD (1.0 µM) and MinE-His (0.75 µM) were prepared together with 30 g/L BSA and 2.5 mM ATP-Mg. For FtsZ reconstitution, FtsZ-Venus-mts (2.5 µM) was prepared with 30 g/L BSA and 50 g/L Ficoll70 as a macromolecular crowder and 2.5 mM GTP-Mg. The co-reconstitution of Min and FtsZ proteins was carried out with the mixture of His-MinD (2 µM), MinE-His (2 µM), His-mScarlet-I-MinC (0.5 µM), and FtsZ-Venus-mts (2.5 µM) together with each 2.5 mM ATP-Mg and GTP-Mg, 30 g/L BSA, and 50 g/L Ficoll70. Confocal images were collected after ~60 min of vesicle preparation, and they were used for up to 1–2 h per sample.

*Cell-free expression inside lipid vesicles*:

Linearized DNA templates were amplified from pPT1-MinD.MinE.Nterm, pPT1-mCherry2-MinC, pPT1-FtsZopt-G55-Venus-Q56, and pPT1-FtsAopt plasmids by using PrimeSTAR Max DNA polymerase (Takara Bio, Shiga, Japan) with T7P-F and T7P-R primers (cccgcgaaattaatacgactcac and caaaaaacccctcaagacccgt), resulting in *min*DE, *mCherry2-min*C, *fts*Z-*Venus*, and *fts*A templates, respectively. Inner solutions for cell-free expression were prepared using PUREfrex 2.0 (GeneFrontier, Chiba, Japan) according to the company's instructions supplied with 10 g/L BSA. The combinations of templates were used for different expression conditions as follows; 0.5 nM *min*DE (for MinDE expression), 3 nM *fts*Z-*Venus* and 1 nM *fts*A (for FtsAZ expression), or 1 nM *min*DE, 1 nM *mCherry2-min*C, 3 nM *fts*Z-*Venus*, and 1 nM *fts*A (for MinCDE/FtsAZ expression). His-msfGFP-MinC was additionally supplied at 0.5 µM to MinDE expression as a fluorescence tracer of MinD localization and 50 g/L Ficoll70 was added to FtsAZ and MinCDE/FtsAZ expression as a crowding agent to enhance FtsZ bundling.

*Microscopy and image processing*:

All confocal images were taken on a Zeiss LSM780 confocal laser scanning microscope using a Zeiss C-Apochromat 20x air (Carl Zeiss). Typically, GUV solution was added to a 384-well plate and placed on LSM780 at room temperature. For temperature control experiments, a homemade chamber was prepared with stacked imaging spacers (Sigma–Aldrich, St. Louis, MO, USA) sandwiched by clean glass coverslips (#1.5, 22 mm × 22 mm, Thermo Fisher Scientific). The glass surface was passivated with 10 g/L BSA solution prior to sample preparation. Chambers were then mounted to the PE120-XY Peltier system (Linkam Scientific Instruments, Surrey, United Kingdom) to control the temperature and placed on LSM780. Membrane (GUVs and SLBs) labeled with Atto 655-DOPE was excited using a 633 nm He-Ne laser, and fluorescence-tagged proteins (EGFP-MinD, mGL-MinD, mScarlet-I-MinC, FtsZ-Venus-mts, and FtsZ-G55-Venus-Q56) were excited using a 488 nm Ar laser or 561 nm laser. Images were typically recorded with a pinhole size of 2.6–4 Airy units at 512 × 512-pixel resolution, either in 3D stacks (typically 13-15 Z-sections with 2-3 µm intervals to visualize whole GUVs membrane surface). The dynamics of Min proteins were further captured by the timelapse recording with 5-20 seconds intervals. To analyze temperature dependency of Min oscillations (Figure 2), the temperature was first raised at 37 ˚C and Min oscillation dynamics was recorded. Then, the temperature was sequentially dropped to 32, 28, 25, and 22 ˚C and at each temperature, Min oscillation dynamics were captured from the identical GUVs.

All recorded images were processed, visualized, and analyzed using Fiji [6] (v1.53 f). Z-stacks were visualized in 3D max or average reconstituted images by the Z projection function. Kymographs were generated using a custom ImageJ macro script described previously [3]. In short, the vesicle periphery was detected from fluorescence signals of either ATTO655-labelled DOPE or GFP and then straightened to obtain 1D line images over the time series. Subsequently, line images were stacked in an orthogonal direction in order of elapsed time to obtain a kymograph. Kymographs were then further analyzed to anlyze the oscillation period of Min waves. The representative section within the kymograph (typically 5 pixels (width) by N pixels (height), where N reflects the number of recorded time points for imaging) was isolated, then scaled to 1 pixel by 10N pixels to obtain 1D line profile of the mGL-MinD intensity over time course at fixed membrane position. This line profile was then fitted to the sine curve function, obtaining the period of Min oscillation within a GUV. To obtain the Ld area coverage, kymographs of membrane intensity were generated as described above based on ATTO655-DOPE fluorescence. The kymographs were then analyzed by using default thresholding function in Fiji to calculate time-averaged Ld coverage on the equatorial sections.

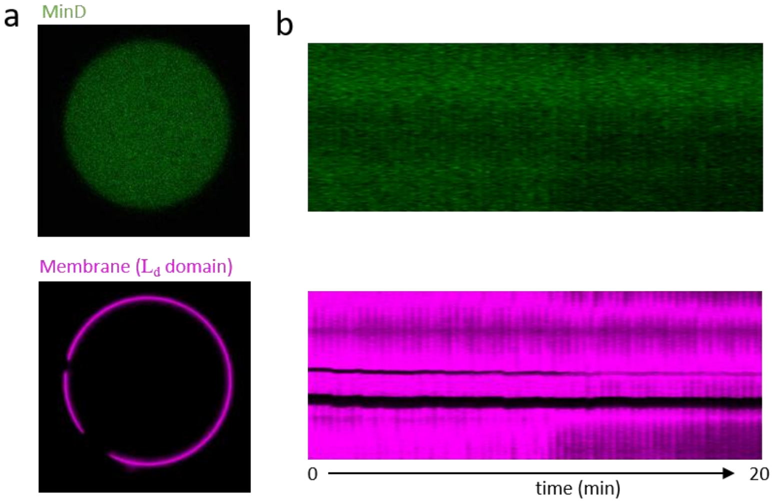


Fig. S1: Phase-separated GUVs composed of DOPC: DPPC: cholesterol without charged lipids exhibit no binding of Min proteins to the membrane. (a) Representative snapshot of a GUV with phase-separated membranes (as shown in $L_d$ domain labelled with Atto-655-DOPE (magenta) and $L_o$ domain is unlabeled (dark)) shows no membrane binding of Min proteins (MinD labelled with EGFP shown in green). (b) Kymograph of membrane localization of Min proteins in a phase-separated GUV shown in (a) confirms that Min proteins exhibit dynamic behaviors neither on low-fluid nor high-fluid domains without charged lipids in the membrane.

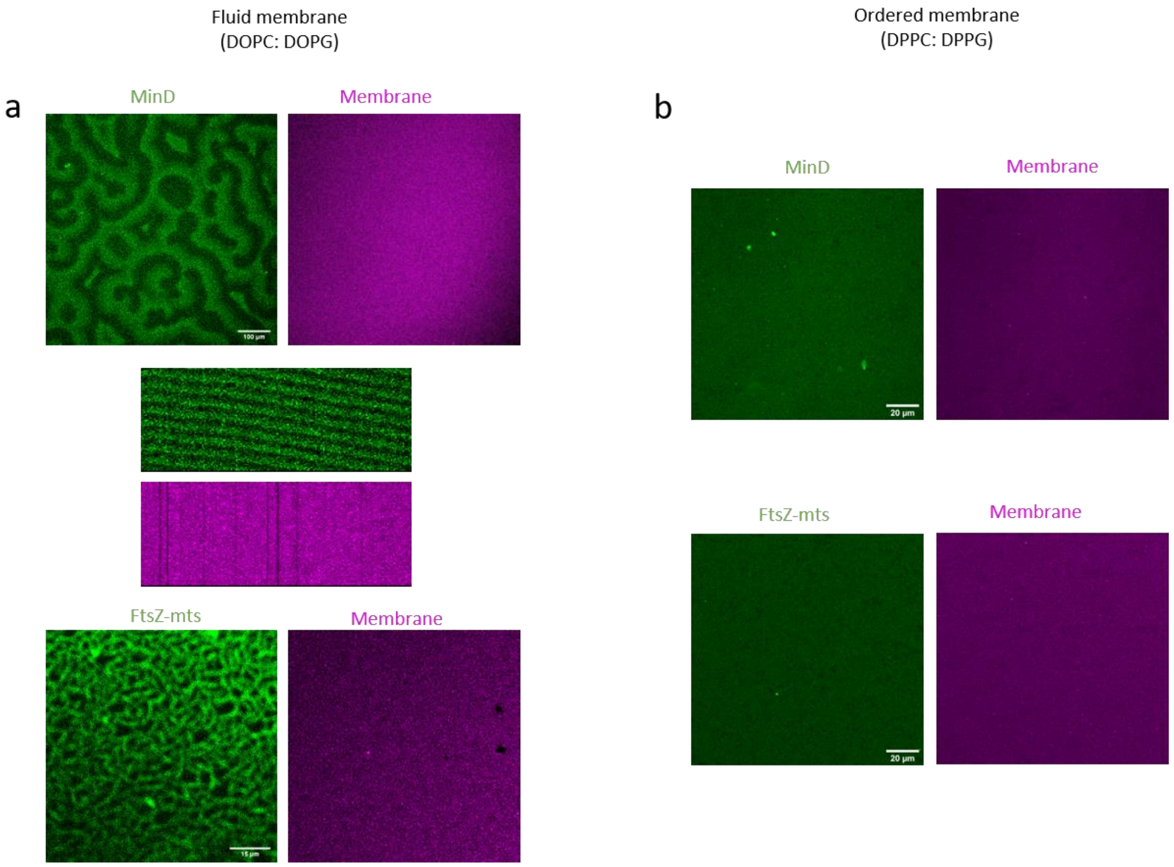


Fig. S2: Self-organization assays of Min/Fts on SLBs of different fluidities. (a) SLBs composed of fluid lipids, i.e. DOPC: DOPG (70: 30) with overall 30 % negative charge labelled with Atto 655 DOPE (magenta) show binding of Min/Fts proteins, where Min proteins (green, top) forms patterns as shown in the kymographs and FtsZ-mts forms filaments (green, bottom). (b) In contrast, SLBs composed of highly ordered lipids, i.e. DPPC: DPPG (70: 30) with overall 30 % negative charge labelled with Atto 647N DPPE (magenta) show no binding of Min/Fts proteins (green) to the membrane, indicating low fluidity of the membrane hinders effective membrane-protein interaction through membrane binding domain of proteins.

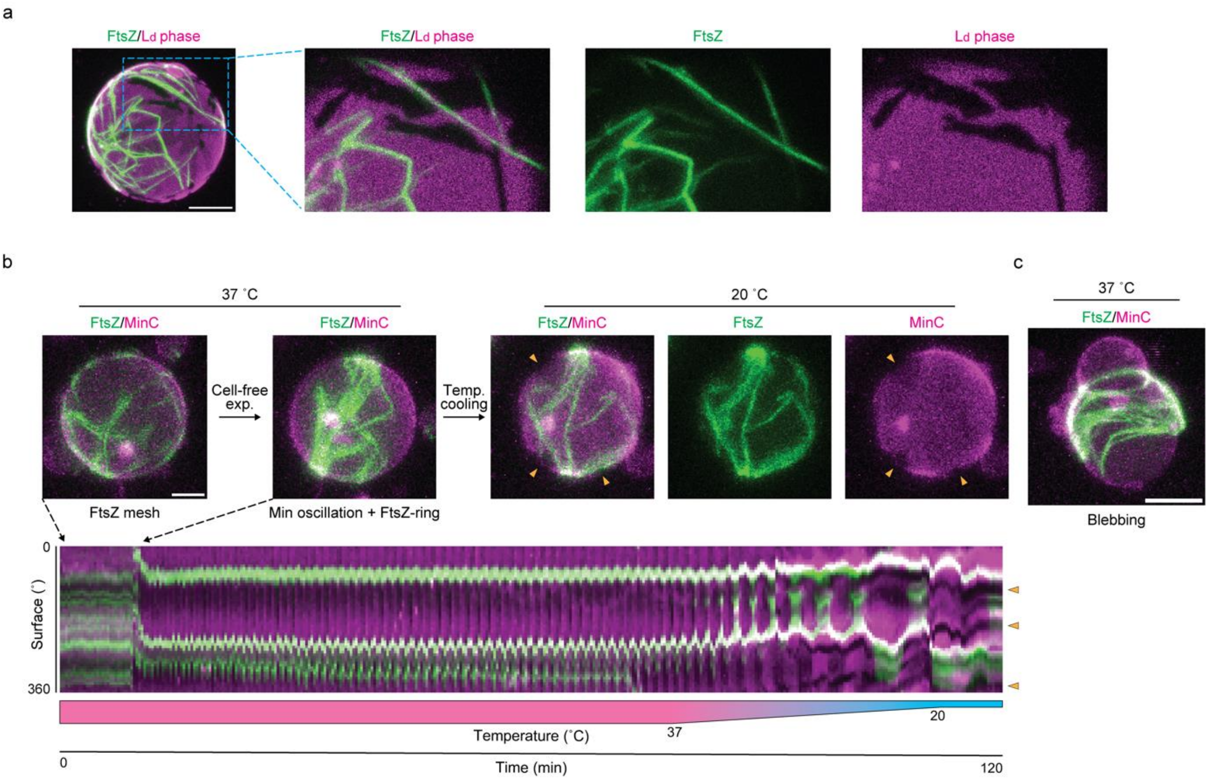


Figure S3: FtsZ-ring formation via cell-free expression of MinCDE and FtsAZ proteins. (a) Cell-free expressed FtsA and FtsZ proteins randomly distributed on the $L_d$ phases within a GUV, confirming that FtsA membrane binding also preferers $L_d$ phases (FtsZ-G55-Venus-Q56 and ATTO655-DOPE are shown in green and magenta, respectively). Scale bar: 10 µm. (b) 3D projection of co-expressed MinCDE and FtsAZ proteins in a phase-separated GUV (top panels) and the kymograph of their dynamics over 120 min period of time (bottom) (FtsZ-G55-Venus-Q56 and mCherry2-MinC are shown in green and magenta, respectively). At the beginning, FtsZ filaments showed random mesh distribution on the membrane, however, once Min oscillation occurred (roughly after 10 min), FtsZ filaments were instantly reorganized into a ring like structure, positioned in the middle of the GUV, and stabilized over 1 h period of time. Furthermore, FtsZ-ring structure remains stable over the temperature cooling process (starts roughly after 1h 20 min) and showed preferential binding to Ld phase together with Min proteins at 20 ˚C. Also see movie S6. Scale bar: 5 µm. (c) Blebbing of the GUV membrane with an internal FtsZ-ring structure surrounding the necking position at 37 ˚C (FtsZ-G55-Venus-Q56 and mCherry2-MinC are shown in green and magenta, respectively). In contrast to the blebbing phenomenon described in Fig. 4c and movie S7, membrane blebbing can be observed without phase-separation, suggesting that membranes show its flexibility above the $T_m$ (Fig. S3c). Scale bar: 10 µm.

#### Supplementary Movies

movie S1: MinDE oscillations in an identical GUV (made from homogeneous membrane condition) with temperature ranging from 37 ˚C to 22 ˚C (mGL-MinD and ATTO655-DOPE are shown in green and magenta, respectively). Oscillation periods becomes significantly slower along with decreasing temperature, showing Arrhenius temperature dependency. Scale bar: 20 µm.

movie S2: Planar projection of the reconstitution of MinCDE/FtsZ in phase separated GUVs (His-mScarlet-I-MinC shown in magenta and FtsZ-mts in green) recorded at 25 ˚C after encapsulation. Min oscillations depict traveling patterns, which sweep the membrane bound FtsZ filaments inside the GUV. Scale bar: 10 µm.

movie S3: 3D projection of the reconstitution of MinCDE/FtsZ in phase separated GUVs (His-mScarlet-I-MinC shown in magenta and FtsZ-mts in green) recorded at 25 ˚C once cycle of heating to 37 ˚C to mix the lipid domains and cooling down to 25 ˚C to yield de-mixed GUVs exhibiting phase separation. After once cycle of temperature sweeps, Min oscillations change from traveling patterns to pole-to-pole which results into FtsZ alignment at the middle of the GUV. Scale bar: 10 µm.

movie S4: 3D projection of Min oscillation via cell-free expressed MinDE proteins in a phase-separated GUV (msfGFP-MinC and ATTO655-DOPE are shown in green and magenta, respectively). MinDE proteins were expressed at 37 ˚C and already indicated oscillation dynamics on the homogeneous membrane in the first 10 min in the movie. Then, as temperature gradually lowered from 37 ˚C to 20 ˚C, phase demixing occurred resulting in membrane "patches", where MinDE preferentially bind to Ld phases (shown in magenta). At the same time, Min oscillation was significantly slowed down along with temperature dropping. Scale bar: 10 µm.

movie S5: 3D projection of Min oscillation via cell-free expressed MinDE proteins in a phase-separated GUV at 20 ˚C (msfGFP-MinC and ATTO655-DOPE are shown in green and magenta, respectively). Min oscillation showed apparent membrane binding to Ld phases (shown in magenta) while showing traveling-like oscillation mode. Scale bar: 5 µm.

movie S6: 3D projection of FtsZ-ring assembly in a phase-separated GUV via cell-free expression of MinCDE and FtsAZ proteins (FtsZ-G55-Venus-Q56 and mCherry2-MinC are shown in green and magenta, respectively). The sequential events of the random mesh distribution of FtsZ filaments, emergence of Min oscillation, and formation of a FtsZ-ring like structure, were recorded at 37 ˚C. After cell-free expression was completed, the temperature cooling process was triggered, and the preferential Ld phase binding of both MinCDE oscillations and the FtsZ-ring structure was confirmed. Scale bar: 5 µm.

movie S7: 3D projection of membrane blebbing processes in a phase-separated GUV with FtsZ-ring after cooling temperature. The GUV already developed FtsZ-ring and Min oscillations via cell-free expression at the beginning of the video (FtsZ-G55-Venus-Q56 and mCherry2-MinC are shown in green and magenta, respectively). After temperature was cooled down until 21 ˚C (roughly after 32 min in the video), the GUV started blebbing from the top-left region where perpendicular and the farthest position from FtsZ-ring. Fascinatingly, the $L_o$ phases (shown as "blank" as neither FtsZ nor MinC binds to $L_o$ regions) were localized on the neck region of the blebbing (also see Figure 4b). The resultant bleb in form of membrane deformation was maintained over 20 min of period after the temperature reached 20 ˚C. Scale bar: 10 µm.
